# Undergraduate Biophysical Chemistry Series: Teaching through a Combination of a Purpose-built Textbook, Research-derived Biomolecular Samples and Computer Labs

**DOI:** 10.64898/2026.08.25.747173

**Authors:** Serge L. Smirnov, Liliya Vugmeyster, Norda Stephenson, James McCarty

**Author notes:** Corresponding Author: Serge L. Smirnov, Jay L. McCarty.

## Abstract

Biophysics is a rapidly advancing field with an incredible breadth of topics. Thus, undergraduate biophysics instructors have to strategize and decide what topics they will cover in their courses. Educational institutions utilize a variety of biophysics textbooks. A common deficiency of each of the existing texts is that it serves well a given set of topics (theory, illustrations, practice problems) and leaves out other areas. A typical example includes good theory and problems for thermodynamics and kinetics while presenting molecular dynamics and various spectroscopic methods in a lacking or outdated way. The authors of this manuscript teach a capstone Biophysical Chemistry three-quarter series (Western Washington University/WWU, Bellingham, WA) which ideally should resonate with the general and major-specific courses the students take within their major at WWU. To achieve this goal and to enrich the traditional lecture-based delivery, the instructors have developed and brought together key pedagogical elements: purpose-built onlinetextbook with a uniform structure of the academic content and practice problems, a study sample (oligopeptide) of biophysical significance with a growing set of experimental and computational data and student-centric in-class activities including computer labs. Our Biophysical series emphasizes concepts and methods of computational structural biology (Molecular Dynamics) and spectroscopic approaches (IR, UV and NMR). Here we describe the details of our integrative approach, summarize key outcomes and chart ways to advance the biophysical chemistry series further. Our textbook can be found through LibreText.

## Introduction

Biophysical Chemistry is an extensive and rapidly evolving field. Knowledge from a number of disciplines underpins modern biophysics, including concepts and skills mastered in general and organic chemistry, physics, biochemistry and a selection of math/calculus courses. Many of these foundational disciplines are well-established and conceptually are changing slowly. For example, the core content of General Chemistry and General Physics have not changed substantially in the last five decades. Contrary to that, modern biophysics includes a number of emerging and rapidly evolving concepts, utilizes an array of progressively more sophisticated experimental and computational methods, and tackles progressively more complex research questions. Because of the integrative nature of the discipline as well as its wide scope and rapid progress, teaching biophysics at an undergraduate level poses several pedagogical challenges. For example, what core principles to cover? What experimental and computational methods to describe (and which ones to leave out)? What types of biophysical research questions to tackle? Whatever the answers to these questions are, the content of the course(s) is typically extensive and challenging.

For these and other reasons, undergraduate biophysical chemistry (biophysics) is often delivered in the form of a capstone, year-long series. In addition to the issues related to the scope of the material to be covered, biophysical chemistry poses other difficulties. The participating students are expected to be able to draw from their prerequisite courses in the areas of multivariable calculus, physics and structural biology of proteins. In reality, students of all levels of comprehension in different disciplines take biophysical chemistry. Thus, the pedagogical infrastructure in this integrative course needs to help students of all levels of capabilities by offering refreshers and practice problems of progressively greater level of sophistication along with the ultimate, most challenging types of questions and exercises. Thus, the pedagogical materials for a biophysics series need to combine a focus on a selected set of concepts, have practice problems covering a range of difficulty levels and have overall structure amenable for student guided and self-directed study during the busy times of academic quarters or semesters. This makes for a very demanding list, which is not easy to find addressed in a commercially available textbook.

To the best of the authors’ knowledge, no currently available biophysics textbook, commercial or free, addresses all these needs and covers all the concepts central to the Biophysics series at WWU at sufficient depth and scope. Commercially available textbooks which provide sufficient learning reinforcement through inter-chapter uniformity of organization and cross-reference of key concepts rarely cover all the areas needed for a specific biophysics course series. This presents a pedagogical and financial disconnect: a text helpful during one quarter can be frustratingly useless in the next one(s). The biophysics series also needs to introduce students to research literature, requiring critical assessment of recent real-world problems from a biophysical perspective. The references in paper textbooks quickly become outdated as the field evolves. Academic concepts and theory are not easy to learn without practical examples. A wet-lab addition is the ultimate way of providing the students with hands-on learning experience. In many cases, adding a wet lab component to a course is not practical due to financial or logistical constraints. Thus, a virtual lab approach (“computer lab”) can be utilized as a way to help the students make connections to the theory and develop the required skills.

Here we present the outcomes of a multi-faceted work our team did to organize and teach a biophysics series at WWU for Biochemistry BS. Our approach combines a free-of-charge online textbook developed by the course instructors equipped with essential pedagogical elements and suitable biomolecular samples with pre-collected experimental and computational data for the virtual/computer labs as well as a set of assessment tools (instruments?) to gauge how well the approach delivers the subject matter and how satisfied the students with the course and their own progression in it.

**Biophysical Chemistry series at Western Washington University** is defined by the academic background of Biochemistry majors and research expertise of the instructors. The three-quarter (CHEM 466, 467, 468) series typically enrolls 32 Biochemistry BS majors during their final / fourth year of undergraduate studies. The class size was somewhat smaller (27 students) during the academic year 2024/25 described here. Typically, the students in this cohort are well-motivated as they are preparing to apply to graduate (PhD or MS) research programs in life sciences as well as to seek employment in pharmaceutical and biotechnology industry. Manageable class size and strong research expertise of the instructors further strengthen the academic environment. Biophysical chemistry aims to expose the students to modern concepts and principles as well as experimental and computational methods utilized in biochemistry, biophysics and related fields. We emphasize concepts and methods of molecular dynamics (MD), various types of spectroscopic methods with emphasis on heteronuclear solution NMR as well as interconnections between these methods.

Each quarter at WWU includes about 10 weeks of instruction time. The first quarter (Fall, Chem 466) of the series is dedicated to the essentials of biochemical thermodynamics and kinetics, statistical mechanics of biomolecules as well as to the foundational principles and methods of molecular dynamics simulations with the focus on structure and dynamics of proteins and peptides. The second quarter (Winter, Chem 467) introduces the students to a number of spectroscopic techniques (e.g., IR, UV/Vis and NMR spectroscopy) and their applications in modern biophysics. The third quarter (Spring, Chem 468) dedicates most of the class time to introducing the students to the process of drafting and composing an original research project proposal in biochemistry or biophysics (broadly defined). The instructional benefits of this multi-week assignment is enhanced by the expectation that in their proposals the students will apply concepts covered in the previous two quarters. The remainder of the Spring quarter typically focuses on select additional topics in biophysics and students participating in the end-of-series survey. Pedagogical infrastructure of the course typically includes a lecture room optimized for students working in small groups, a computer lab and availability of resources to generate samples and data (computational and experimental) for the study samples relevant for ongoing biophysical research. Issues and limitations of the series included the absence of a suitable textbook (as described above), disparity in the students’ academic background and their extra-curricular experiences (engagement in academic research vs. no research). The work presented in this manuscript outlines the measures we took to address the identified academic and organizational challenges while capitalizing on the strengths including (a) design of a custom online textbook emphasizing undergraduate pedagogy, (b) generation of research-inspired biomolecular test samples and (c) synergistically integrating concepts covered in lectures and problem-solving sessions with skills developed in hands-on computer labs across the series.

## Results

### Biophysical Chemistry Textbook: pedagogical principles and content structure

The two instructors (SLS, JM) developed a freely accessible, web-based textbook for the Biophysical Chemistry series (CHEM 466, 467). Designed for senior B.S. Biochemistry majors, the textbook features pedagogical elements beneficial for undergraduate education: (1) uniform structure and organization of every chapter including learning objectives, chapter text and figures as well as examples and practice problems; (2) optimized content covering both traditional biochemical thermodynamics and modern spectroscopic methods. We deliberately designed the content to be minimally sufficient for the course without unnecessary details which may hinder students reading the text. While a prototype of this text has been used in our teaching since Fall 2021, the online format allows regular updates and revisions. Ongoing improvements include integrating the instructor’s research activities into the text, adding more practice problems, and illustrative examples as well as fixing typos and expanding on some chapters. At the time of writing this article, the text consists of six parts (1–6), each including several chapters (4–6). Overall, the text includes 24 chapters, 76 Figures & Tables, 105 examples and problems. All the artwork and problems are original products. Chapters conclude with practice problems with solutions provided separately to encourage the students to attempt solving the problems on their own first. The online format enables students to easily cross-reference material from previous chapters. The textbook is published with LibreText.org under CC BY-NC-SA 4.0 license: https://chem.libretexts.org/Courses/Western_Washington_University/Biophysical_Chemistry_(Smirnov_and_McCarty)

Some chapters from parts 5 and 6 were also used for teaching undergraduate and graduate biophysics at University of Colorado / Denver, e.g. in Structural Biology (undergraduate/MS level) and Graduate Biochemistry classes during Spring 2022 and Fall 2025.

### A Research-derived oligopeptide case study

To introduce students to modern and experimental and computational methods in biophysics, we utilize the research-derived oligopeptide Gly_1_Gly_2_Lys_3_Gly_4_Met_5_Gly_6_Phe_7_Gly_8_Leu_9_ (**Figure 1A**). The molecule is denoted RC9 (nine-residue random coil peptide) and was chosen because of its anticipated disordered state.

**Figure 1.**
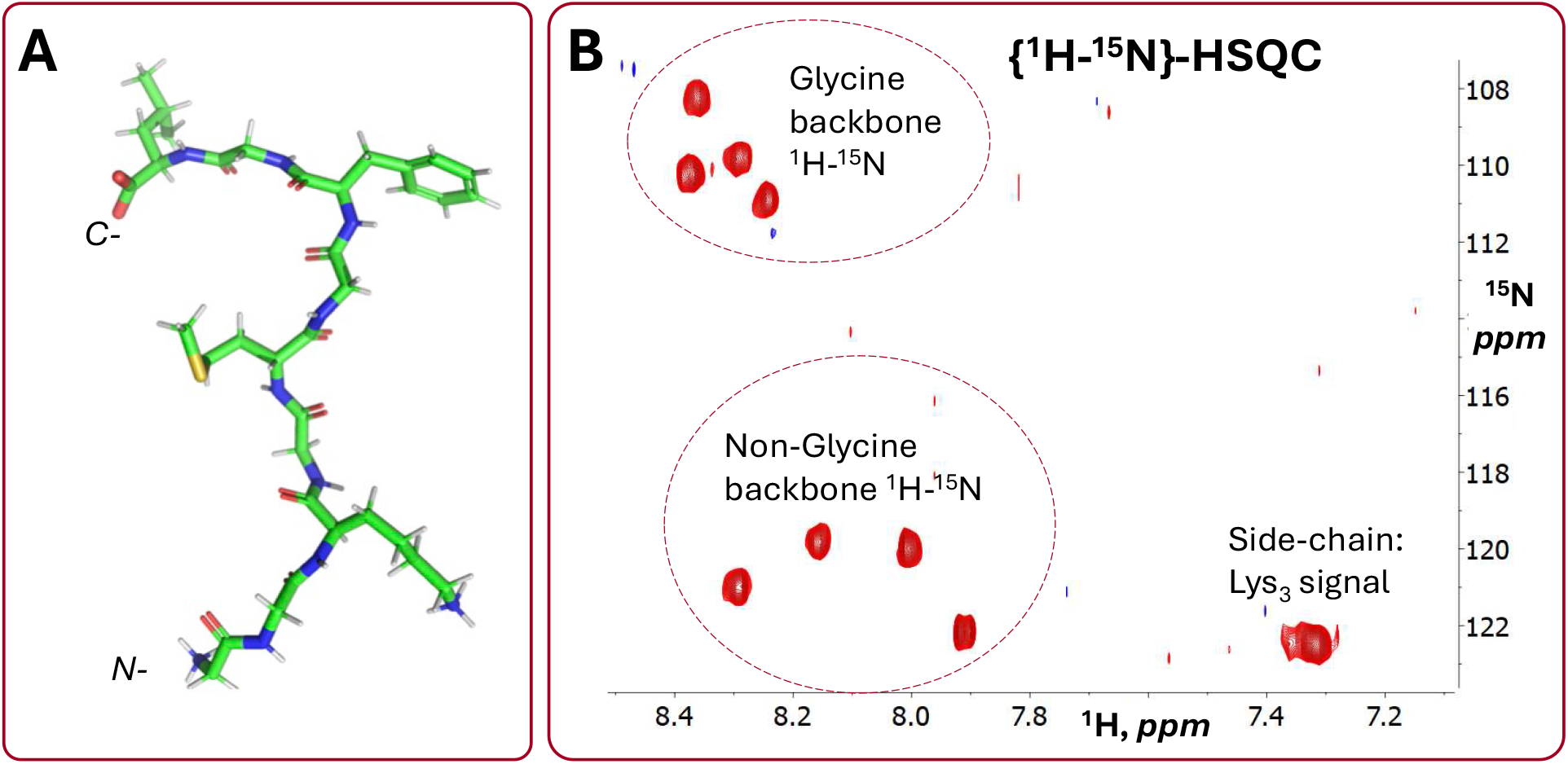
**A**: RC9 peptide, GGKGMGFGL, used as a case study in Biophysical Chemistry series. The amino and carboxy termini of the molecule are labeled N- and C- respectively. **B**: Solution {^1^H-^15^N}-HSQC NMR spectrum of 2.0 mM RC9. The spectrum was recorded at ^15^N natural abundance, pH 2, with a Bruker Avance III HD spectrometer equipped with a room-temperature TXI probe.

RC9 oligopeptide is an excellent model compound for developing solution and solid-state NMR relaxation methodologies to probe backbone and side-chain dynamics via incorporation of site-specific isotopic labels, including studies using quadrupolar nuclei such as ^2^H and ^17^O in addition to more commonly used ^13^C and ^15^N isotopes.^1-2^ Further, due to its disordered nature it can be used for comparison of dynamics with the disordered regions of functional proteins, such as flexible unstructured regions of plant villins, amyloid-β and tau proteins.^3-5^ RC9 was characterized by solution NMR and CD spectroscopy and is being used for further solid-state NMR studies (**Figure 1B**).^2, 6^ These spectra and resonance assignments offer a rich set of opportunities to introduce the key types of NMR spectra and to demonstrate how these data are collected and analyzed for a comprehensive structural studies of protein-like samples. The instructors of the WWU Biophysical Chemistry course also performed 1-microsecond molecular dynamics (MD) simulations for this same RC9 sample, with and without conformational restraints, as described in the Materials & Methods section.

“Big Ideas” help to integrate Computer Labs with Lectures and Testing

The instructors tried to augment their traditional lecturing with active student work in small groups. In addition, the computer labs were developed to visualize and analyze molecular dynamics trajectories of the target polypeptides (CHEM 466) as well as to analyze structure and dynamics implications of solution NMR spectra for the same samples (CHEM 467). In both cases, the computer labs were structured around a set of progressively more sophisticated and interconnected big ideas.

The MD computer lab Big Ideas (an 80-min session in CHEM 466):

- **Big Idea 1**: Equilibrium is the statistical balance of dynamical events. The RC9 peptide is an example, but this idea applies to any biomolecule. The MD simulation shows the peptide rapidly interconverts between several possible configurations (just like a butane molecule interconverts between eclipsed, gauche, and trans configurations). The individual molecule wiggles and jiggles in response to the force exerted on each atom by all the surrounding atoms. If MD simulation trajectory is long enough so that each state is visited many times, the fraction of time spent in each state is related to the equilibrium **probability** to be in that state, and this probability is given by the Boltzmann distribution.
- **Big Idea 2**: Even though the RC9 peptide is constantly fluctuating, some transient conformations are energetically favorable because of relatively weak non-covalent interactions (like hydrogen bonds, salt bridges, and cation-pi interactions that form and break over time). These stabilizing, non-covalent interactions shift the probability in favor of some conformations more than others, shaping the ensemble of structures observed in the MD simulation trajectory.
- **Big Idea 3**: At equilibrium, configurations from MD simulations are determined from the underlying free energy landscape according to the Boltzmann factor. This determines the sampling probability. By working backwards, we can use the observed sampled distributions to *infer* this underlying free energy surface from a histogram of observed states. In this way, we can reconstruct the free energy surface by using the frequency of sampled states because this sampling reflects the equilibrium distribution.

The Computer Lab activity for CHEM 466 was designed to provide hands-on experience for students to interpret a 1-*μs* MD trajectory of RC9 oligopeptide. Students visualized the molecular trajectory, assessed the RC9 structure and dynamics, calculated several observables, and related the MD trajectory to the histogram of observables. Two lectures had been delivered prior to the computer lab, covering the basics of intermolecular interactions, potential energy surfaces, and molecular dynamics. A pre-lab survey was administered via Canvas and was due before the computer lab to assess student’s prior knowledge. The MD computer lab consisted of three parts:

1. Students use PyMOL (The PyMOL Molecular Graphics System, Version 3.0 Schrödinger, LLC) to visualize the 1-*μs* MD trajectory of RC9, visually identifying extended and compact configurations (**Figure 2A)**
2. Students use the MDAnalysis^7-8^ library in Python to compute the end-to-end distance (distance between *C_α_* of residues Gly1 and Leu9), the Root Mean Square Deviation (RMSD) relative to a fully extend chain, and the radius of gyration (Rg). Student calculate the mean value, standard deviation, and a histogram of the distribution (**Figure 2B)**. Python analysis codes were provided as shared Jupyter notebooks, using the cloud-based environment of Google Colaboratory (Google, 2024).
3. In Part 3, students use Chemiscope,^9^ an interactive structure/property explorer for materials and molecules, to interactively explore how individual structural conformations from the MD simulation map to points on a two-dimensional scatter plot of Rg vs. RMSD, along with their associated probability (**Figure 2C**).

**Figure 2.**
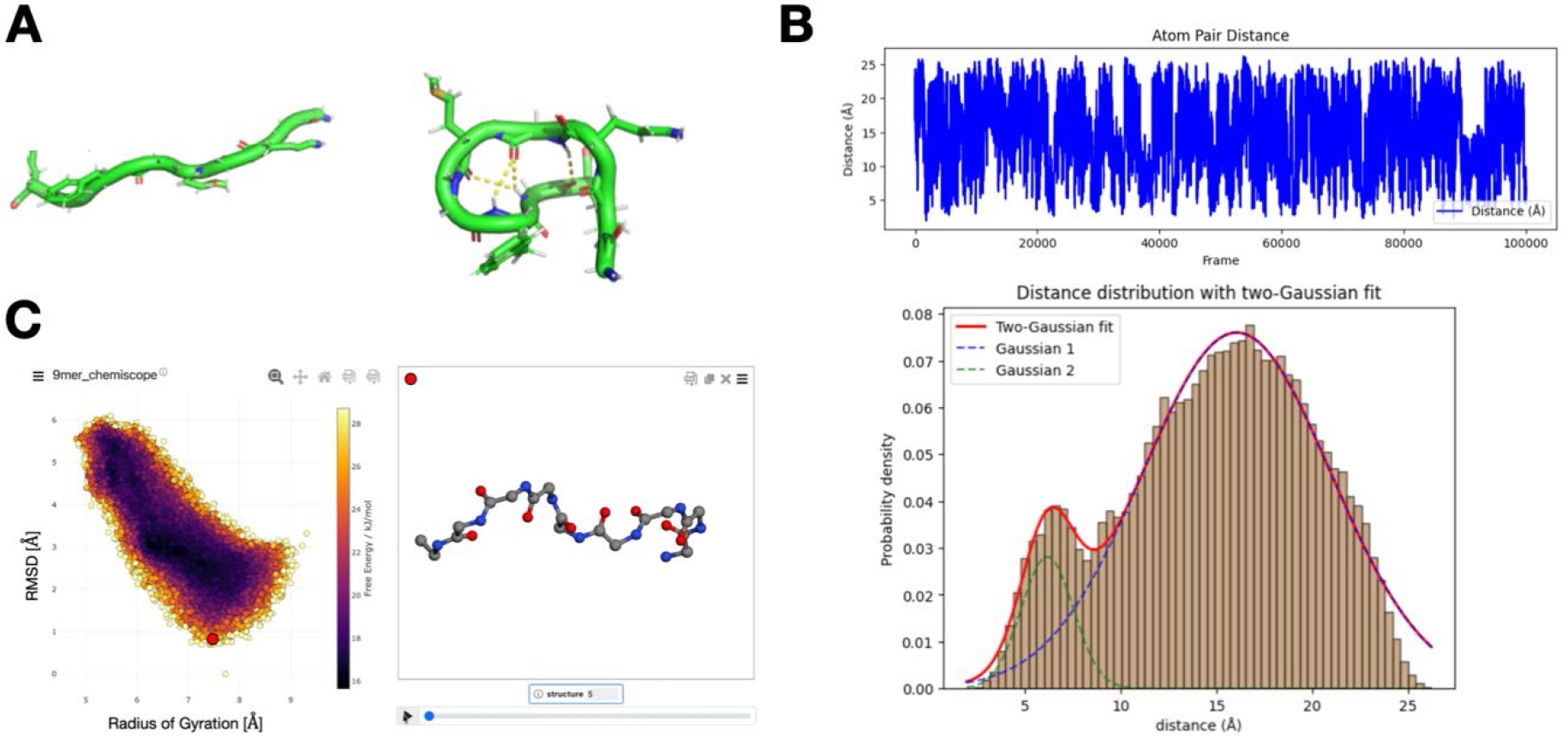
**A**: Students use PyMOL to visualize extended and compact configurations of RC9 peptide from a 1-*μs* MD trajectory. **B**: Students use the Python to compute structural properties and histograms from the MD simulation data. **C**: Students view an interactive Chemiscope representation that shows how individual structures from the MD trajectory map onto points within a scatter plot of RMSD vs. Radius of Gyration.

After completing the in-class portion of the computer lab, students receive a take-home post-lab assignment to be completed as graded homework. In this follow-up assignment, students complete a worksheet with embedded links to additional MD simulation trajectories and Python Jupyter notebooks. Students are asked to compare structural properties from the 1-*μs* MD trajectory of RC9 oligopeptide to those from a 36 *μs* MD simulation of the folding of the 10-residue chignolin mini-protein, GYDPETGTWG, performed by DE Shaw group (**Figure 3A**).^10^ Finally, students investigate liquid structure by computing the radial distribution function from a provided MD simulation trajectory of liquid argon, based on Rahman’s original 1964 paper.^11^ The provided Python Juypter notebook guides students through the numerical Fourier transform and comparison to experimental scattering function (**Figure 3B).**

**Figure 3.**
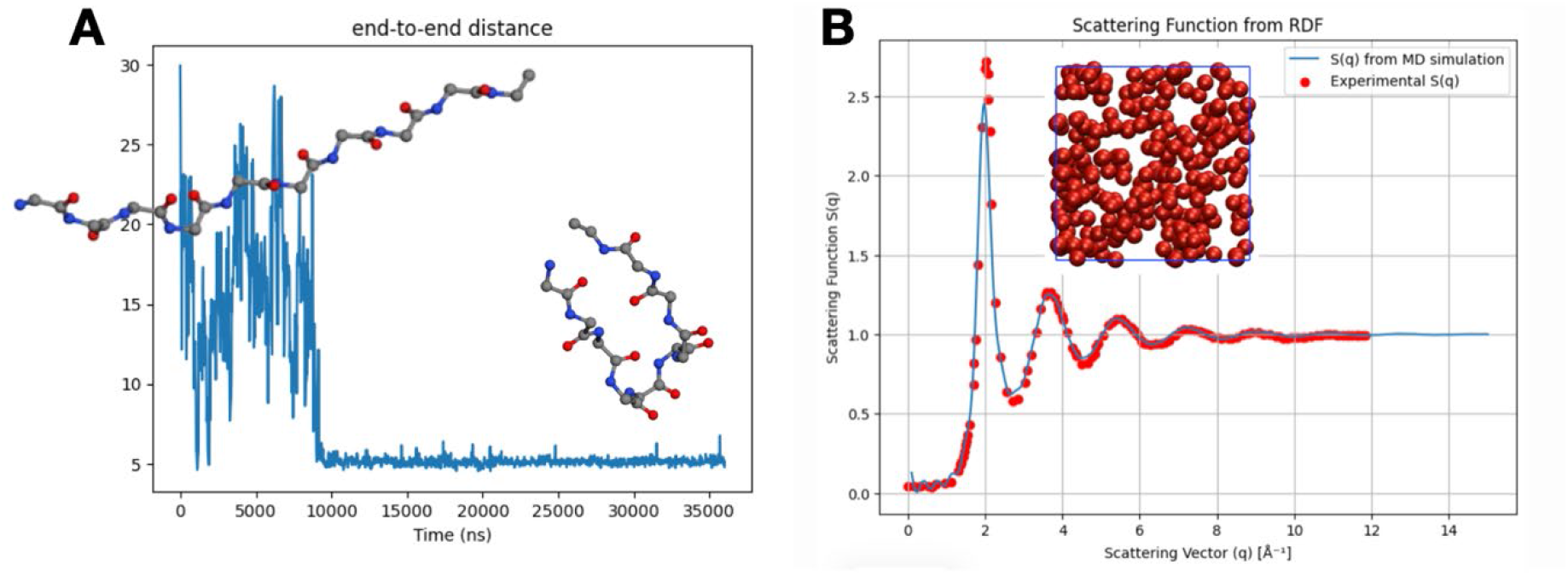
**A**: Students visualize the end-to-end distance and representative structures of the 10- residue chignolin mini-protein performed from D.E. Shaw. **B**: Students use Python to compute the scattering function for liquid argon and compare to experimental data.

Heteronuclear solution NMR computer lab Big Ideas (two 50-min sessions in CHEM 467):

- **Big Idea 1**: Equilibrium is the statistical balance of dynamic events and structural heteronuclear solution NMR spectra, including 2D {^1^H-^13^C}-HSQC and {^1^H-^15^N}- HSQC, report signals representative of the statistical weighted average of the sample conformations. The RC9 and polypeptide samples of similar size offer simple and clean spectra, which are easy to interpret and which nevertheless have most properties of typical spectra recorded for full-size proteins.
- **Big Idea 2**: Structural homeo- and hetero- nuclear NMR spectra, including 2D {^1^H-^13^C}-HSQC, {^1^H-^15^N}-HSQC, {^1^H-^1^H}-NOESY and {^1^H-^1^H}-TOCSY, offer a coherent set of data sufficient to perform the residues-specific NMR resonance assignment. The resonance assignment process helps the students to build connections between their perception of the primary structure (amino acid sequence) of the sample, the nature of the specific type of NMR data (e.g., each signal in {^1^H-^15^N}-HSQC represents a covalent ^1^H-^15^N group, **Figure 1**) and online repositories of the NMR chemical shifts (e.g., Biomolecular Magnetic Resonance databank, BMRB entry 51754 for RC9).^6, 12^
- **Big Idea 3**: A combination of the residues-specific NMR resonance assignments and 2D {^1^H-^1^H}-NOESY signal strength values offer a set of structural distance and dihedral values restraints for robust 2°/3° structure characterization of a polypeptide sample. The instructors share with the students first-hand knowledge of converting the NMR data into biomolecular structure and into dynamic characterization of intrinsically disordered proteins.^13-17^

After the lab session, the follow-up take-home exercise include a digital simulation of the Fourier transform of the raw NMR signal (electric current recorded vs. time) to generate an NMR spectrum (spectral intensity vs. resonance frequency), prediction of the NMR spectra of selected types (e.g., 2D {^1^H-^1^H}-TOCSY and {^1^H-^15^N}-HSQC) expected for RC9 oligopeptide, comparison of the student-predicted spectra of these types with actual spectra distributed at the end of the lab session as well as performing the resonance assignment of NMR signals for a target polypeptide sample, e.g. RC9 (e.g., combining published resonance assignments from BMRB entry 51754 and newly acquired NMR data, **Figures 1, 4**). Students submit these outcomes for grading as post-lab reports. In addition to the post-lab reports, in Chem 466 and 467 graded homeworks and exams were administered to allow the instructors to assess students’ comprehension of the material. In both quarters, the practice problems discussed in class were typically fairly basic, homework questions added another level of complexity and the take-home exams which followed offered a broad range of question difficulty, from simple to advanced.

**Figure 4.**
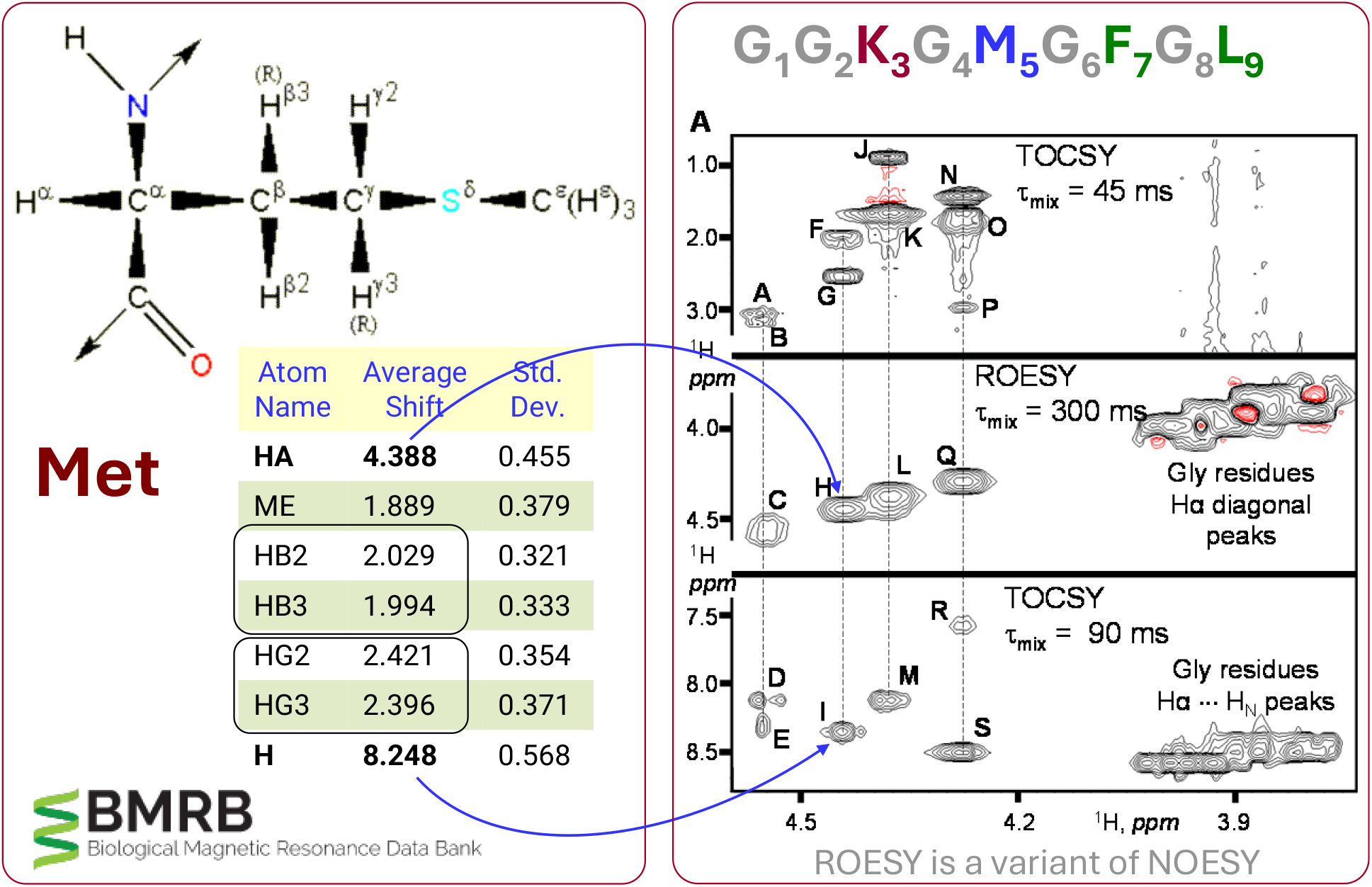
A general set of concepts and data used to perform the ^1^H NMR resonance assignment of the RC9 peptide sample. Central types of ^1^H NMR data used for that include 2D ^1^H NOESY (ROESY) and 2D ^1^H TOCSY, each collected with at least two different mixing time values. For the assignment of certain ^13^C and ^15^N resonances, respective 2D heteronuclear spectra ({^1^H-^13^C}- HSQC and {^1^H-^13^C}-HSQC) are used (Figure 1).

### Assessment of Learning Outcomes through Surveys

The students were offered several groups of questions to probe their retained knowledge and self-assessment in the subject matter together with objective testing of their knowledge. These were grouped into numerical self-assessment responses as well as multiple-choice and open-answer test-like questions. For the most part, this activity was graded on completion to lower stress anxiety and encourage broader participation. Many questions listed in **Tables 1**, **2** and **3** below are mean to assess student learning of the Big Ideas (listed above) as well as conceptual foundation and technical elements of the computational and experimental methods covered in CHEM 466/467. These assessments were administered at five time points throughout the series: before and after the practical computer labs in Chem 466 and Chem 467 (pre-lab and post-lab respectively) as well as at the end of the Spring quarter (Chem 468).

**Table 1.** Numerical Self-Assessment Questions and Outcomes.

| **CHEM 466 (Fall quarter)** | **Questions** | **Average±Std.Dev**<br>(Pre-Lab) → (Post-Lab) | **Cohen's d** |
| --- | --- | --- | --- |
| **1.T1_466** | At what level can you explain the basics of a molecular dynamics (MD) simulation algorithm at the level of being able to outline its steps? | (1.7 ± 0.8) → (2.7 ± 0.8) | 1.33 |
| **3.T1_466** | At what level can you describe the Boltzmann distribution? | (2.4 ± 0.6) → (3.1 ± 0.8) | 1.04 |
| **#4.T1_466** | At what level can you apply the Boltzmann distribution to equilibrium systems? | (2.2 ± 0.7) → (2.7 ± 0.7) | 0.75 |
| **6.T1_466** | At what level can you describe the relationship between free energy, probability, and sampling of conformational states? | (2.4 ± 0.8) → (3.3 ± 0.8) | 1.15 |
| **8.T1_466** | At what level can you relate the inter-atomic distance distributions from a molecular dynamics trajectory to the free energy? | (1.5 ± 0.6) → (2.7 ± 0.6) | 2.00 |
| **CHEM 467 (Winter quarter)** | **Questions** | **Average±Std.Dev**<br>(Pre-Lab) → (Post-Lab) | **Cohen's d** |
| **1.T1_467** | At what level can you explain the basics of the NMR effect for spin-$\frac{1}{2}$ particles? | (1.5 ± 0.5) → (3.0 ± 0.7) | 2.47 |
| **#3.T1_467** | At what level can you apply the Boltzmann distribution to systems at equilibrium? | (2.6 ± 0.7) → (3.1 ± 0.6) | 0.77 |
| **4.T1_467** | At what level can you explain how any two types of NMR data can be plugged into a molecular dynamics simulation to determine the 3$^{\circ}$ structure of a | (1.8 ± 0.5) → (2.6 ± 0.7) | 1.32 |
| **5.T1_467** | At what level can you estimate inter-proton distances utilizing a 2D NOESY NMR spectrum with both dimensions assigned to specific hydrogens? | (1.0 ± 0.2) → (2.6 ± 0.8) | 2.74 |

\*Question identifiers in this and all the next tables include question number (before the period “.”
and reference to the specific table and course). E.g., **3.T1\_466** refers to question 3 in Table 1, section describing CHEM 466. In these self-assessment questions, the students ranked their ability to answer each question as “not able”, “slightly able”, “moderately able” and “extremely able” quantified as 1, 2, 3 and 4 respectively in the “Average ± Std.Dev.” column.

**Table 2.** Multiple-choice Assessment Questions and Outcomes.

| CHEM 466 (Fall quarter) | Multiple-Choice Questions and Outcomes | Average±Std.Dev<br>(Pre-Lab) → (Post-Lab) | Cohen's d |
| --- | --- | --- | --- |
| #1.T2_466 | What experimental technique(s) can be used to report a protein tertiary structure? (select all that apply): (A) X-ray crystallography; (B) solution NMR spectroscopy; (C) Circular Dichroism spectroscopy; (D) Molecular Dynamics simulation; (E) I do not know. | (3.0 ± 0.8) → (3.3 ± 0.6) | 0.41 |
| 3.T2_466 | In a molecular dynamics simulation, Newton's equation of motion is solved: (A) numerically and iteratively; (B) numerically and only once to determine the motion; (C) analytically and iteratively; (D) analytically and only once to determine the motion; (E) Newton's equation of motion is not used at all in molecular dynamics; (F) I do not know. | (2.9 ± 1.3) → (3.4 ± 1.1) | 0.40 |
| 7.T2_466 | How does the molecular dynamics trajectory resemble an experiment? (A) A molecular dynamics trajectory reproduces the true time evolution of the real system; (B) Time averages computed from the molecular dynamics trajectory represent ensemble averages from an experiment; (C) A molecular dynamics trajectory has no real physical basis and therefore no connection to experiment; (D) Both A and B; (E) I do not know. | (2.8 ± 1.3) → (3.6 ± 0.7) | 0.82 |
| 8.T2_466 | How does the free energy landscape affect the sampling of conformational states in an MD simulation? (A) It has no effect since the simulation is already at equilibrium; (B) It determines how frequently different conformational states are sampled; (C) It only affects the initial part of the simulation until equilibrium is reached, but does not affect the equilibrium distribution; (D) It is negligible because the system has enough thermal kinetic energy at 310 K to sample all conformational states; (E) I do not know. | (2.5 ± 1.1) → (3.6 ± 0.9) | 0.93 |
| #9.T2_466 | How do you think MD simulations can be used to interpret or complement solution NMR 3<sup>o</sup> structure/dynamics data? (A) By directly predicting chemical shifts; (B) By providing static distances for a single conformation; (C) By providing time-averaged structures and distance distributions; (D) By simulating the effect of the nuclear spin angular moment in a magnetic field; (E) I do not know. | (2.7 ± 1.3) → (3.8 ± 0.7) | 1.03 |
| CHEM 467 (Winter quarter) | Multiple-Choice Questions and Outcomes | Average±Std.Dev<br>(Pre-Lab) → (Post-Lab) | Cohen's d |
| #1.T2_467 | What experimental technique(s) can be used to determine a protein 3O structure? (select all that apply): (A) X-ray crystallography; (B) solution NMR spectroscopy; (C) Circular Dichroism spectroscopy; (D) Molecular Dynamics simulation; (E) I do not know. | (3.0 ± 0.8) → (3.4 ± 0.6) | 0.58 |
| 2.T2_467 | In a molecular dynamics simulation, Newton's equation of motion is solved: (A) numerically and iteratively; (B) numerically and only once to determine the motion; (C) analytically and iteratively; (D) analytically and only once to determine the motion; (E) Newton's equation of motion is not used at all in molecular dynamics; (F) I do not know. | (3.5 ± 1.1) → (3.4 ± 1.1) | -0.11 |
| 3.T2_467 | Which types of inter-atomic interactions in proteins can be assessed with the help of a suitable solution NMR experiment (Select all that apply): (A) van der Waals interactions between closely positioned hydrogens; (B) a short network of covalent bonds connecting two hydrogen atoms; (C) backbone dihedral angle values; (D) hydrogen bonds; (E) electrostatic interactions; (F) I do not know. | (2.4 ± 0.8) → (3.0 ± 0.9) | 0.75 |
| #4.T2_467 | How do you think MD simulations can be used to interpret or complement solution NMR 3<sup>o</sup> structure/dynamics data?: (A) By directly predicting chemical shifts; (B) By providing static distances for a single conformation; (C) By providing time-averaged structures and distance distributions; (D) By simulating the effect of the nuclear spin angular moment in a magnetic field; (E) I do not know. | (3.2 ± 1.2) → (3.4 ± 0.9) | 0.22 |

\*For each question, a rubric was developed to grade the answers uniformly. The respective numerical scores ranged from 1 (poor answer or no answer) to 4 (excellent answer).
#These questions were asked in both CHEM 466 and Chem 467.

**Table 3.** Open-ended Assessment Questions and Outcomes.

| **CHEM 466 (Fall quarter)** | **Open-ended Questions** | **Average±Std.Dev**<br>(Pre-Lab) → (Post-Lab) | **Cohen's d** |
| --- | --- | --- | --- |
| **4.T3_466** | In your understanding, what are some thermodynamic, tertiary structure, or dynamic properties that might be computed from MD simulations? | (2.2 ± 0.7) → (2.8 ± 0.7) | 0.85 |
| **5.T3_466** | Consider an intrinsically disordered protein that can adopt several distinct conformational states. In your own words, describe how the instantaneous value of the backbone dihedral angles relates to the observed experimental values of these angles? | (2.4 ± 1.0) → (2.7 ± 0.9) | 0.35 |
| **CHEM 467 (Winter quarter)** | **Open-ended Questions** | **Average±Std.Dev**<br>(Pre-Lab) → (Post-Lab) | **Cohen's d** |
| **4.T3_467** | In your understanding, what are some thermodynamic properties that might be obtained from heteronuclear solution NMR spectra? | (1.4 ± 0.7) → (2.0 ± 0.9) | 0.83 |
| **5.T3_467** | In your understanding, what are some tertiary structure properties that might be obtained from heteronuclear solution NMR spectra? | (1.4 ± 0.6) → (2.5 ± 0.8) | 1.58 |
| **6.T3_467** | In your understanding, what are some backbone dynamic properties that might be obtained from heteronuclear solution NMR spectra? | (1.3 ± 0.6) → (2.0 ± 1.1) | 0.78 |
| **7.T3_467** | For an intrinsically disordered protein, describe how a heteronuclear solution NMR spectrum of this protein may differ from the one recorded from a folded protein of similar size? What type of NMR spectrum would be useful for such a comparison? | (1.6 ± 0.7) → (2.7 ± 1.1) | 1.19 |

\*For each question, a rubric was developed to grade the answers uniformly. The respective numerical scores ranged from 1 (poor answer or no answer) to 4 (excellent answer).

### Numerical Responses for Students’ Self-Assessment

Student’s self-assessment is discussed in this section with a set of select questions for Chem 466 and Chem 467 (**Table 1**). Some of these select questions and their pedagogical purposes are discussed below. Each question offers four options for self-rating: 1. Not able at all; 2. Slightly able; 3. Moderately able; and 4. Extremely able. For each question, the table reports the class average ± standard deviation and the “Cohen’s d” comparison of the two stages during the quarter: at the start (before the practical computer lab or pre-lab) and at the end of the quarter (after the lab, Post-Lab). For Chem 467, the pre-lab self-assessments were collected during weeks 2 and 3 whereas the post-lab took place after the computer lab organized during week 8. For Chem 466, the pre-lab self-assessments were collected during week 8 after students had been introduced to statistical mechanics of biomolecules in lectures but prior to the computer lab activity. The MD computer lab activity occurred during week 9 and the post-assessment was collected during week 11 (Finals Week).

The select questions allow the students to rate their understanding of foundational concepts related to the molecular dynamics as a computational method and NMR spectroscopy of spin-½ systems as an experimental approach and their applications for structural biology of biomolecular samples. The purpose and outcomes for some of these questions are discussed here. For both Chem 466 and Chem 467, questions **1**.T1_466 and **1**.T1_467 are asking the students to rate their grasp of the foundations, the molecular dynamics algorithm and NMR effect for spin-½ nuclei respectively. Questions **2**.T1_466 and **3**.T1_466 as well as question **3**.T1_467 are asking the students to rate their fluency with the central idea and applications of the Boltzmann distribution. This concept is discussed as a central concept for both classes, which helps enhance the students’ awareness and knowledge retention to fairly high levels (class average rose from 2.2 to 3.1). While similar, Questions **3**.T1_466 and **4**.T1_466 target different levels of student cognitive ability. Specifically, Questions **3**.T1_466 asks about student’s ability to explain/describe, whereas Question **4**.T1_466 asks student’s to specifically assess their ability to apply this knowledge. Question **6**.T1_466 asks about students’ ability to connect multiple concepts: equilibrium, conformational sampling, probability, and free energy. Question **8**.T1_466 is probing the students’ confidence in relating the set of frames forming the MD trajectory with a distribution of one of the most basic features- interatomic distances. Question **4**.T1_467 represents an extension of this idea: it requests to self-reflect on what types of structural NMR data (inter-atomic distances, dihedrals, etc.) can be incorporated into a molecular dynamics simulation to convert purely computation-based “free MD” into an experiment-based “restrained MD” simulation. Furthermore, question **5**.T1_467 is asking to think about one’s capacity to interpret 2D NOESY NMR data and how it can be used to extract distance restraints from the spectral properties (e.g., NOE signal intensities). Many questions in this self-assessment survey are probing the students’ confidence regarding the Big Ideas underpinning the computer labs. Specifically, questions **4**.T1_466 and **5**.T1_466 target learning outcomes related to Big Idea 1-3 for Chem 466 (the connection between dynamic equilibrium, Boltzmann probability, and free energy), and questions **1**.T1_467 and **3**.T1_467 are linked with Big Idea 1 for Chem 467 lab (“structural” solution NMR spectra are reporting signal patterns corresponding to the ensemble-averaged conformations of the polypeptide). Questions **4**.T1_467 and **5**.T1_467 are cumulatively probing students’ grasp of Big Ideas 2 and 3: NMR resonance assignment and structure determination.

The pedagogical effect on student’s self-assessment was quantified with “**Cohen’s d**” metric (**Table 1**) with absolute values below 0.2 generally represent a small effect, over 0.5 generally described as indicating medium effect and 0.8 or greater – large effect.^18^ All the questions except **4**.T1_466 (Chem 466) and **3**.T1_467 (Chem 467) had Cohen’s d value greater than 1.04 reliably indicating large positive pedagogical effects of our teaching approach which combines lectures, textbook and computer labs. Questions #4 (Chem 466) and #3 (Chem 467) had Cohen’s d value of 0.75 and 0.77 respectively, which correspond to a medium effect and approach the large effect bracket (threshold 0.80). The relatively smaller scale of improvement for these two questions could be attributed to the fact that the concept of Boltzmann distribution is one of the most abstract ideas in computational and experimental structural biology and that it does take a sequence of two quarters to advance the student confidence in a major way (from 2.2 to 3.1 for the class average). This relatively high gain in confidence is very much in line with “*Repetitio est mater studiorum*”, a classic Latin proverb which values repetition as the mother of studying.

### Student Multiple-Choice Answers

This section describes multiple-choice questions which were designed to test the students’ grasp of key concepts. In this section, multiple answers were possible for each question to reflect the complexity of the concepts studied. Numerical scoring of the answers was done as per a set of rules to reflect all meaningful combinations of selected options.

**Table 2** lists representative questions probing the students’ grasp of central concepts introduced in CHEM 466 and 467. The question style in many cases resembles that of those found in exams and homeworks during the respective quarters. The questions in this table represent probing of the students’ prior knowledge (e.g., questions 1.T2_466, 2.T2_466), their grasp of the new material specific to each class (e.g., questions 3.T2_466, 3.T2_467) and connections between the two quarters (questions 9.T2_466, 4.T2_467). Generally, Cohen’s d values show a medium to large positive effect of our teaching (five out of nine questions have Cohen’s d value exceeding 0.5, **Table 2**).

We noted that for the questions in which the students had little experience (lower class average scores in the pre-lab part of the table 2) their learning gains were more substantial (Cohen’s d >0.5). If the class had significant level of prior knowledge, the learning gains were more modest (**Figure 5**). For CHEM 466, the relatively high starting (pre-lab) average values for questions **1**.T2_466 and **3**.T2_466 reflect substantial prior knowledge available to the students. Material relevant for question **1**.T2_466 was covered to a certain degree in prerequisite Biochemistry courses and concepts central for 3.T2_466 were introduced in CHEM 466 itself prior the survey. Although the numerical magnitude of teaching effect is relatively low for these questions (Cohen’s d value of ∼0.4), the post-lab class average values reflect robust grasp of the material (class average ≥3.3). Teaching during the next quarter of the series (CHEM 467) builds on the foundation established in CHEM 466. Thus, some questions are repeated in the part of Table 2 for CHEM 467 although not all the related concepts are explicitly taught during this quarter. In CHEM 467 part of Table 2, two out of four questions (1.T2_467 and 3.T2_467) showed a noticeable improvement (Cohen’s d > 0.5). The subject of question 2.T2_467 was not taught in CHEM 467 but the question was asked in Winter quarter to assess the retention of this important piece of knowledge over the winter break. Analysis of identical questions yielded no significant change in the outcomes for questions 3.T2_466 (post-lab average of 3.4) and 2.T2_467 (post-lab average of 3.4). During CHEM 467 the variation of the class average, although negative, is numerically insignificant (3.5 to 3.4 with Cohen’s d of 0.11). We conclude that the students were able to retain the knowledge related to these questions over the duration of one quarter when these concepts were not explicitly talked about by the instructor. Question 4.T2_467 shows a noticeable drop in class average which emerged after the winter break. The class average value for this question improved post-lab in CHEM 467 but it remained somewhat lower than at the end of the previous quarter (CHEM 466) which may be explained by several nuanced reasons outlined later in the paper.

**Figure 5.**
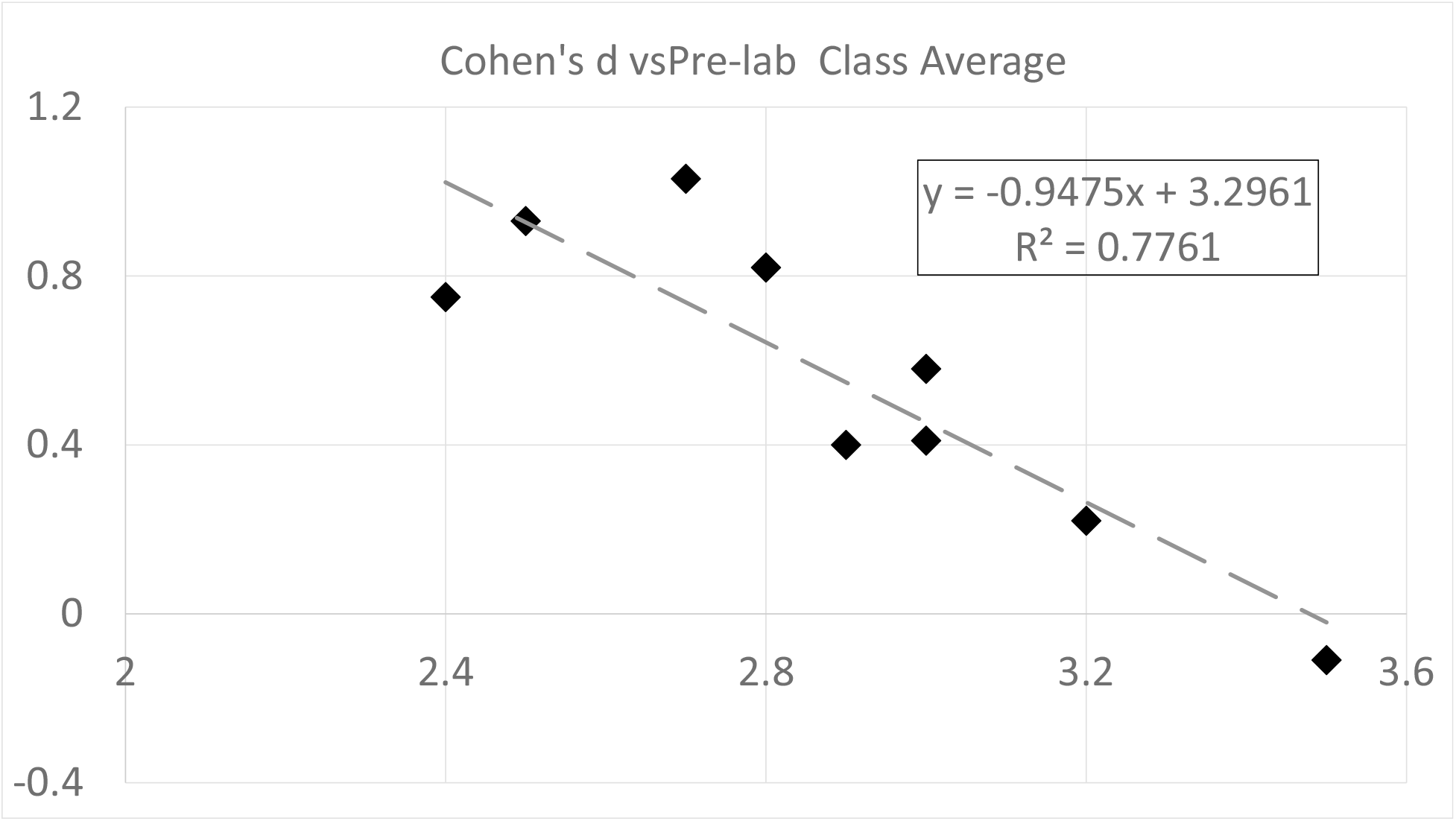
Analysis of gains in knowledge as a function of prior knowledge state based on data in Table 2. Cohen’s d value shows negative correlation with the of class pre-lab knowledge (class average).

### Open-ended free-response questions

Open-ended questions were designed to assess the students’ grasp of the new concepts and their capacity and willingness to utilize new terms introduced during the respective quarters of the biophysics series.

The open-ended questions were added to the survey to probe the capacity of the students to develop and formulate thoughts on the subject matter utilizing the terms introduced during the two starting quarters of the series (**Table 3**). The questions presented in **Table 3** invite students to present their thoughts on how the two central methods of the quarters (MD simulations for CHEM 466 and heteronuclear solution NMR spectroscopy for CHEM 467) can be utilized to study thermodynamic, structural and dynamic properties of biomolecules. The standard types of samples in both quarters were proteins, both folded and intrinsically disordered.

Answers to all the questions indicated large positive effects of the combined implementation of our in-class lectures and student group work as well as the textbook and computer labs (Cohen’s d at 0.8 or greater in most cases). The utilization of the new terms introduced during CHEM 466 and 467 noticeably improved in the target cohort.

### End-of-series student assessment: how much do the students forget?

During the third quarter of the series (Chem 468, Spring) the main student’s activity was to prepare a research proposal in the area of biomolecular sciences. No new major Biophysical Chemistry concepts were covered and no systematic reviews of those introduced and taught in Fall/Winter were conducted. However, the instructor of CHEM 468 encouraged the students to apply the knowledge of the methods discussed in the previous quarters of the series to be applied for their research proposal. The learning outcomes during the Fall (Chem 466) and Winter (Chem 467) quarters were generally encouraging and positive (**Tables 1**, **2** and **3**). Thus, we were eager to utilize similar sets of questions to assess the knowledge retention by the students over longer time (three months or longer) when no reviews of the previously taught concepts were offered.

Some representative questions probing knowledge retention by the end of the Biophysical Chemistry series are presented in **Table 4**. The assessment done in the Spring quarter shows that the students mostly retained their knowledge and confidence in applying these concepts: Cohen’s d values were either positive or rarely below ™0.5, which corresponds to minor to moderate negative effect of time on memory and knowledge.

**Table 4.** Knowledge retention assessed over time periods exceeding one academic quarter.

| Numerical Self-Assessment Questions and Outcomes (subset from Table 1) | Average±Std.Dev | Cohen's d |
| --- | --- | --- |
|  | (CHEM 466 Post-Lab) → (CHEM 468) |  |
| 1.T1_466 At what level can you explain the basics of a molecular dynamics (MD) simulation algorithm at the level of being able to outline its steps? | (2.7 ± 0.7) → (2.7 ± 0.8) | 0.05 |
| 3.T1_466 At what level can you describe the Boltzmann distribution? | (3.1 ± 0.8) → (3.1 ± 0.7) | 0.06 |
| 4.T1_466 At what level can you apply the Boltzmann distribution to equilibrium systems? | (2.7 ± 0.7) → (2.8 ± 0.7) | 0.17 |
| 6.T1_466 At what level can you describe the relationship between free energy, probability, and sampling of conformational states? | (3.3 ± 0.8) → (3.2 ± 0.7) | -0.11 |
| 8.T1_466 At what level can you relate the inter-atomic distance distributions from a molecular dynamics trajectory to the free energy? | (2.7 ± 0.6) → (2.8 ± 0.8) | 0.05 |
|  | (CHEM 467 Post-Lab) → (CHEM 468) |  |
| 1.T1_467 At what level can you explain the basics of the NMR effect for spin-½ particles? | (3.0 ± 0.7) → (3.2 ± 0.8) | 0.27 |
| 2.T1_467 At what level can you explain the difference between a homonuclear 2D NMR experiment and a heteronuclear one? | (3.5 ± 0.6) → (3.4 ± 0.8) | -0.17 |
| 3.T1_467 At what level can you apply the Boltzmann distribution to systems at equilibrium? | (3.1 ± 0.6) → (2.8 ± 0.7) | -0.46 |
| 4.T1_467 At what level can you explain how any two types of NMR data can be plugged into a molecular dynamics simulation to determine the 3° structure of a protein? | (2.6 ± 0.7) → (2.7 ± 0.8) | 0.09 |
| 6.T1_467 At what level can you explain how the three key parameters of recorded raw NMR signal (initial current intensity, frequency of the current oscillations and the rate of the signal relaxation) relate to the three key properties of a related NMR spectral signal (intensity or height, peak position and line width at half-height)? | (3.2 ± 0.7) → (3.1 ± 0.7) | -0.17 |
| 7.T1_467 At what level can you explain the differences in the NMR R1 and R2 relaxation rates for folded/ordered vs. unfolded/disordered polypeptides? | (2.6 ± 0.9) → (2.7 ± 0.9) | 0.07 |
| Multiple-Choice Assessment Questions and Outcomes (subset from Table 2) | Average±Std.Dev | Cohen's d |
|  | (CHEM 466 Post-Lab) → (CHEM 468) |  |
| 1.T2_466 What experimental technique(s) can be used to report a protein tertiary structure? (select all that apply): (A). X-ray crystallography; (B). solution NMR spectroscopy; (C). Circular Dichroism spectroscopy; (D). Molecular Dynamics simulation; (E). I do not know. | (3.4 ± 0.5) → (3.4 ± 0.6) | 0.00 |
| 3.T2_466 In a molecular dynamics simulation, Newton's equation of motion is solved: (A) numerically and iteratively; (B) numerically and only once to determine the motion; (C) analytically and iteratively; (D) analytically and only once to determine the motion; (E) Newton's equation of motion is not used at all in molecular dynamics; (F) I do not know. | (3.5 ± 0.9) → (3.9 ± 0.5) | 0.49 |
| 7.T2_466 How does the molecular dynamics trajectory resemble an experiment? (A) A molecular dynamics trajectory reproduces the true time evolution of the real system; (B) Time averages computed from the molecular dynamics trajectory represent ensemble averages from an experiment; (C) A molecular dynamics trajectory has no real physical basis and therefore no connection to experiment; (D) Both A and B; (E) I do not know. | (3.6 ± 0.8) → (3.5 ± 0.7) | -0.20 |
| 8.T2_466 How does the free energy landscape affect the sampling of conformational states in an MD simulation? (A) It has no effect since the simulation is already at equilibrium; (B) It determines how frequently different conformational states are sampled; (C) It only affects the initial part of the simulation until equilibrium is reached, but does not affect the equilibrium distribution; (D) It is negligible because the system has enough thermal kinetic energy at 310 K to sample all conformational states; (E) I do not know. | (3.7 ± 0.8) → (3.5 ± 1.1) | -0.22 |
|  | (CHEM 467 Post-Lab) → (CHEM 468) |  |
| 1.T2_467 What experimental technique(s) can be used to determine a protein 3O structure? (select all that apply): (A) X-ray crystallography; (B) solution NMR spectroscopy; (C) Circular Dichroism spectroscopy; (D) Molecular Dynamics simulation; (E) I do not know. | (3.4 ± 0.6) → (3.4 ± 0.6) | 0.07 |
| Open-ended Assessment Questions and Outcomes (subset from Table 3) | Average±Std.Dev | Cohen's d |
|  | (CHEM 466 Post-Lab) → (CHEM 468) |  |
| 2.T3_466 What is a PDB file and what information does it have? | (2.9 ± 0.9) → (2.9 ± 0.9) | 0.00 |
| 4.T3_466 In your understanding, what are some thermodynamic, tertiary structure, or dynamic properties that might be computed from MD simulations? | (2.8 ± 0.7) → (2.6 ± 0.7) | -0.22 |
|  | (CHEM 467 Post-Lab) → (CHEM 468) |  |
| 4.T3_467 In your understanding, what are some thermodynamic properties that might be obtained from heteronuclear solution NMR spectra? | (2.0 ± 0.9) → (2.4 ± 1.4) | 0.34 |
| 5.T3_467 In your understanding, what are some tertiary structure properties that might be obtained from heteronuclear solution NMR spectra? | (2.5 ± 0.8) → (3.1 ± 0.9) | 0.72 |
| 7.T3_467 For an intrinsically disordered protein, describe how a heteronuclear solution NMR spectrum of this protein may differ from the one recorded from a folded protein of similar size? What type of NMR spectrum would be useful for such a comparison? | (2.7 ± 1.1) → (3.0 ± 1.2) | 0.30 |

Some questions were deliberately repeated across the series to assess the relative effects of instructors’ teaching and students’ decline of competency over time of infrequent usage of the newly learned terms and concepts. A good example is the question probing the students’ confidence in their ability to apply the Boltzmann distribution to a macroscopic system at equilibrium (questions **4**.T1_466, **3**.T1_467, **3**.T1_468). For this question, the class average went from 2.2 (Chem 466, pre-lab) to 3.1 (Chem 467, post-lab) with Cohen’s d of 1.38 indicating a strong positive effect of teaching. This major improvement in students’ confidence came about in spite of a small decrease in the class average value between the end of the Fall (class average 2.7) and start of the Winter quarters (class average 2.6). As expected, student self-confidence in answering this question regressed by the end of the Spring quarter during which the related concepts were not formally discussed: the class average went from 3.1 (Chem 467, post-lab) to 2.8 (Chem 468) with Cohen’s d of ™0.46 (small to medium effect). Overall, our teaching was beneficial as the class average increased from 2.2 (pre-lab, Fall quarter) to 2.8 (end of the Spring quarter) and Cohen’s d value of 0.87, which corresponds to a strong positive effect.

Another example is open-ended Question 4.T3_466 (Table 3), which assesses higher-order thinking by asking students to consider potential applications of MD simulations across thermodynamics, structure, and dynamics. The class average score increased from 2.2 in the pre-lab survey to 2.8 immediately following the in-class MD activity (Table 3, Cohen’s d of 0.85; large effect). This improvement is consistent with students having just completed an activity where they calculated several properties from the MD trajectory of RC9 nonapeptide. The average score decreased slightly to 2.6 on the follow-up survey administered in CHEM 468 (five months after the in-class MD activity). The decline represents only a small effect (Cohen’s d of ™0.22). The overall improvement from the pre-survey in CHEM 466 to the follow-up survey had a medium effect size (Cohen’s d of 0.63), suggesting students retain an improved ability and leave the three-quarter series able to identify structural, thermodynamic, and dynamic properties that can be computed from MD simulations. A similarly positive dynamic was observed for a related question probing the students’ grasp of what protein structural properties can be inferred from heteronuclear solution NMR spectra of such samples (question 5.T3_467 and 5.T3_468). The overall improvement was noticeable during the Winter CHEM 467 quarter (Cohen’s d value of 1.58, **Table 3**) and at the end of the Spring CHEM 468 quarter (positive Cohen’s d of 0.72, **Table 4**). Related questions (4.T3_467 and 6.T3_467) assessed during the Winter and Spring quarters showed more modest but still robust improvements overall (**Table 4**). No loss of class proficiency for the NMR-related questions was registered for the NMR-themed questions after three months at the end of the Spring CHEM 468 quarter (all Cohen’s d values are positive).

Considering the Multiple-Choice Question, **3**.T2_466 (**Table 2**), students appear to continue to improve after CHEM 466 post-assessment survey (**Table 4**). The class average score increased from 3.4 all the way to 3.9 in CHEM 468. (Cohen’s d of 0.49 is a small-medium effect). This is likely a result of additional lectures and homework presented in the final weeks of CHEM 466, and students growing familiarity with biophysical methods across the series. By the end of the series, nearly all the students correctly understand that MD simulation involves iteratively solving Newton’s equation numerically on the computer to generate a discrete time trajectory.

## Materials and Methods

### RC9 Molecular Dynamics simulations

1-microsecond molecular dynamics (MD) simulations for RC9 oligopeptide were run with and without conformational restraints. All-atom molecular dynamics (MD) simulations of RC9 in water were performed using GROMACS 2019.4 with the CHARMM36m protein force fields and TIP3P water model. The starting structure was a fully extended chain built using PyMOL. The system was solvated in a cubic box with a volume of 195.112 nm^3^ with periodic boundary conditions. The system was equilibrated for 100 ps in the NVT ensemble at 300 K using the V-rescale thermostat, followed by an NPT equilibration in the NPT ensemble for 1000 ps at 300 K and 1 bar using a Parrinello-Rahman barostat. We generated a 1 *μ*s-long continuous MD trajectory of RC9 without constraints and a separate 1 *μ*s-long continuous MD trajectory with a harmonic restraint between atom HA1 on Gly2 and atom HN on LEU9. This restrained limits the sampling of the extended conformations, yielding a structurally distinct ensemble for students to analyze.

### RC9 heteronuclear solution NMR spectroscopy

A range of homonuclear and heteronuclear NMR spectra were acquired for this sample including 1D ^1^H collected at different temperature values as well as homonuclear 2D ^1^H COSY, TOCSY, NOESY, ROESY and heteronuclear {^1^H- ^13^C}-HSQC and {^1^H-^15^N}-HSQC (**Figure 1**) utilizing unlabeled (natural abundance) and site-specifically labeled samples (^13^C labeling and deuteration of the side chain methyl group of the central methionine). The ^1^H/^13^C NMR resonance assignments for the sample are reported publicly with Biological Magnetic Resonance Data Bank (BMRB entry 51754). To visualize and analyze the 1D and 2D NMR spectra, the students used MestReNova software, version [17.0.1-41952], Mestrelab Research S.L., Santiago de Compostela, Spain (http://www.mestrelab.com).

### Assessment Surveys

The assessment surveys (**Tables 1**, **2**, **3**, **4**) were administered via Canvas at five points during the progression of the series: two before and after the computer labs in Chem 466 (Fall quarter) and Chem 467 (Winter quarter) and at the end of Chem 468 (Spring quarter).

## Discussion

The assessment data analyzed here indicates that the students’ learning outcomes were generally positive throughout the Biophysical Chemistry series. The class average improvements during each specific quarter were positive. Moreover, our assessment approach went beyond a single- quarter time frame and allowed to probe longer-term knowledge retention spanning months after key concepts were covered in class. At the end of the series, knowledge retention was shown to be generally robust while simultaneously learning new concepts and effects of students’ forgetting the new material small (**Tables 4**).

A single outlier in this positive trend is presented by question **9**.T2_466 (**Table 2**) where considerable gains in knowledge made in Chem 466 were partially lost during the inter-quarter period (Winter break) and the loss was not fully recovered during Chem 467. Although the class average change and Cohen’s d metric indicated overall improvement between the start and end points of the series, the observed partial loss of students’ comprehension between the quarters will inform the instructor of Chem 467 to update their teaching of the relevant concepts. This will be achieved by identifying and addressing common misconceptions, enhancing the computer lab and dedicating more class time to the concepts which can be aided by greater number of credit hours assigned to Chem 467 starting Winter 2027.

Suggestions and ideas for the next iterations of the series: create a uniform for all three quarters master list of questions (not every question will be asked during every quarter). The intent here is to create a question bank to streamline and enhance assessment in Chem 468. Some questions will be rephrased to increase their clarity and ease of answer interpretation (why the students choose specific answers). A greater number of MC questions will be matched to respective open-ended and self-assessment questions to solicit more nuanced responses and better interpret the self-assessment responses. Chem 467 will utilize a greater class time as it goes from 3 contact hours in Winter 2026 to 4 contact hours starting Winter 2027. The extra time will allow to address some concepts deeper and to broaden the material covered. New study samples (oligopeptides) will be generated for their computational (MD) and experimental (NMR) characterization as well as for the introduction of this data into the textbook. Respective enhancements to the computer lab will be added in Chem 466 and Chem 467.

## Conclusion

A three-quarter Biophysical Chemistry series at WWU is now equipped with a purpose-built textbook, advanced study samples derived from research activities of the instructors, an advanced computer-lab components all utilized in a student-centered environment. Students gain direct experience with research-derived data for the RC9 oligopeptide system, emphasizing the two-way synergy between computational and heteronuclear NMR methods for studying protein conformational dynamics. The data presented and discussed here provides a foundation for further improvements including novel study samples, updates of the free-of-charge online textbook, and a refined master assessment set of questions. All these developments are expected to further enhance student learning and long-term knowledge retention by clarifying challenging concepts in biomolecular statistical physics.

## AUTHOR INFORMATION

### Author Contributions

JM and SLS: conceptualized the work, collected experimental and computational data, developed pedagogical materials, developed & implemented assessment tools, analyzed the assessment data, drafted and finalized the manuscript.

LV: developed experimental samples and collected initial data with the samples, drafted and proofread significant segments of the manuscript.

NS: developed assessment tools, analyzed the assessment data, drafted and proofread the manuscript.

### Funding Sources

JM was funded by the NSF (CAREER award CHE-2102189). NS funding TBA. The work was supported by an award from WWU /RSP to SLS and JM to design the Biophysical Chemistry textbook. Initial RC9 sample design and generation was supported by LV.

## ACKNOWLEDGMENT

We thank …

## Notes

https://chem.libretexts.org/Courses/Western_Washington_University/Biophysical_Chemistry_(Smirnov_and_McCarty)

### Competing Interest Statement

The authors have declared no competing interest.

## REFERENCES

1. Vugmeyster, L.; Ostrovsky, D.; Rodgers, A.; Gwin, K.; Smirnov, S. L.; McKnight, C. J.; Fu, R., Persistence of Methionine Side Chain Mobility at Low Temperatures in a Nine-Residue Low Complexity Peptide, as Probed by (2) H Solid-State NMR. Chemphyschem 2024, 25 (4), e202300565.

2. Vugmeyster, L.; Ostrovsky, D.; Fu, R., Carbon-detected deuterium solid-state NMR rotating frame relaxation measurements for protein methyl groups under magic angle spinning. Solid State Nucl Magn Reson 2024, 130, 101922.

3. Clarke, S.; Pedersen, W.; Heins, J.; Howerton, C. M.; Curry, E. J.; Mosier, D. L.; Pillitteri, L. J.; Antos, J. M.; Smirnov, S. L., Actin filament bundling by the Arabidopsis VILLIN4 C-terminus involves the basic segment of the IDR linker and is sensitive to temperature and ionic strength. under review with PLoS ONE 2026.

4. Au, D. F.; Ostrovsky, D.; Fu, R.; Vugmeyster, L., Solid-state NMR reveals a comprehensive view of the dynamics of the flexible, disordered N-terminal domain of amyloid-beta fibrils. J Biol Chem 2019, 294 (15), 5840–5853.

5. Zhang, J. Y.; Dregni, A. J.; Hong, M., Heterogeneous Dynamics of the Fuzzy Coat of Full-Length Phospho-Mimetic Tau Fibrils. J Am Chem Soc 2026, 148 (1), 1623–1637.

6. Vugmeyster, L.; Ostrovsky, D.; Rodgers, A.; Gwin, K.; Smirnov, S. L.; McKnight, C. J.; Fu, R., Persistence of Methionine Side Chain Mobility at Low Temperatures in a Nine-Residue Low Complexity Peptide, as Probed by 2H Solid-State NMR. Chemphyschem 2024, 25(*4*) (Epub 2024 Jan 4.).

7. Gowers, R. J.; Linke, M.; Barnoud, J.; Reddy, T. J. E.; Melo, M. N.; Seyler, S. L.; Dotson, D. L.; Domanski, J.; Buchoux, S.; Kenney, I. M.; O., B., MDAnalysis: A Python package for the rapid analysis of molecular dynamics simulations. Proceedings of the 15th Python in Science Conference 2016, 98–105.

8. Michaud-Agrawal, N.; Denning, E. J.; Woolf, T. B.; Beckstein, O., MDAnalysis: a toolkit for the analysis of molecular dynamics simulations. J Comput Chem 2011, 32 (10), 2319–27.

9. Fraux, G.; Cersonsky, R. K.; Ceriotti, M., Chemiscope: Interactive Structure-Property Explorer for Materials and Molecules. Journal of Open Source Software 2020, 5 (51), 2117.

10. Lindorff-Larsen, K.; Piana, S.; Dror, R. O.; Shaw, D. E., How fast-folding proteins fold. Science 2011, 334 (6055), 517–20.

11. Rahman, A., Correlations in the motion of atoms in liquid argon. Physical review 1964, 136.2A, A405.

12. Romero, P. R.; Kobayashi, N.; Wedell, J. R.; Baskaran, K.; Iwata, T.; Yokochi, M.; Maziuk, D.; Yao, H.; Fujiwara, T.; Kurusu, G.; Ulrich, E. L.; Hoch, J. C.; Markley, J. L., BioMagResBank (BMRB) as a Resource for Structural Biology. Methods Mol Biol 2020, 2112, 187–218.

13. Miears, H. L.; Gruber, D. R.; Horvath, N. M.; Antos, J. M.; Young, J.; Sigurjonsson, J. P.; Klem, M. L.; Rosenkranz, E. A.; Okon, M.; McKnight, C. J.; Vugmeyster, L.; Smirnov, S. L., Plant Villin Headpiece Domain Demonstrates a Novel Surface Charge Pattern and High Affinity for F-Actin. Biochemistry 2018, 57 (11), 1690–1701.

14. Gruber, D. R.; Toner, J. J.; Miears, H. L.; Shernyukov, A. V.; Kiryutin, A. S.; Lomzov, A. A.; Endutkin, A. V.; Grin, I. R.; Petrova, D. V.; Kupryushkin, M. S.; Yurkovskaya, A. V.; Johnson, E. C.; Okon, M.; Bagryanskaya, E. G.; Zharkov, D. O.; Smirnov, S. L., Oxidative damage to epigenetically methylated sites affects DNA stability, dynamics and enzymatic demethylation. Nucleic Acids Res 2018, 46 (20), 10827–10839.

15. Fedechkin, S. O.; Brockerman, J.; Pfaff, D. A.; Burns, L.; Webb, T.; Nelson, A.; Zhang, F.; Sabantsev, A. V.; Melnikov, A. S.; McKnight, C. J.; Smirnov, S. L., Gelsolin-like activation of villin: calcium sensitivity of the long helix in domain 6. Biochemistry 2013, 52 (45), 7890–900.

16. Fedechkin, S. O.; Brockerman, J.; Luna, E. J.; Lobanov, M. Y.; Galzitskaya, O. V.; Smirnov, S. L., An N-terminal, 830-residue Intrinsically Disordered Region of the Cytoskeleton-regulatory Protein Supervillin Contains Myosin II- and F-actin- Binding Sites. J Biomol Struct Dyn 2013, 31(*10*), 1150–1159.

17. Boyko, K. V.; Rosenkranz, E. A.; Smith, D. M.; Miears, H. L.; Oueld Es Cheikh, M.; Lund, M. Z.; Young, J. C.; Reardon, P. N.; Okon, M.; Smirnov, S. L.; Antos, J. M., Sortase-mediated segmental labeling: A method for segmental assignment of intrinsically disordered regions in proteins. PLoS One 2021, 16 (10), e0258531.

18. Hedges, L. V., Interpretation of the Standardized Mean Difference Effect Size When Distributions Are Not Normal or Homoscedastic. Educ Psychol Meas 2025, 85 (2), 245–257.

